# Peritubular macrophages and spermatogonia are sequentially increased in the testis of rats after mono-(2-ethylhexyl) phthalate exposure

**DOI:** 10.1101/767707

**Authors:** Ross Gillette, Richa Tiwary, Jorine JLP Voss, Shavini N Hewage, John H Richburg

**Author notes:** **AUTHOR CONTRIBUTIONS** Conceptualization: J.H.R., R.T., J.J.L.P.V.; Methodology: R.G., R.T., J.W.V.; Investigation: R.G., R.T., J.J.L.P.V., S.N.H.; Formal Analysis: R.G.; Writing – Original Draft: R.G.; Writing – Review & Editing: R.G., R.T., J.H.R.; Funding Acquisition: J.H.R. **Declaration of Interest:** The authors declare no competing interests. These authors contributed equally to the work (Ross Gillette, Richa Tiwary).

## Abstract

Peripubertal exposure to the phthalate metabolite mono-(2-ethylhexyl) phthalate (MEHP) in rodents causes testicular inflammation, spermatocyte apoptosis, and disruption of the blood-testis barrier. The MEHP-induced inflammation response includes an infiltration of macrophages and neutrophils to the testes, although the cause and purpose of this response is unknown. Recently, a population of testicular macrophages phenotypically distinct from those resident in the interstitium was described in mice. Testicular peritubular macrophages aggregate near the spermatogonial stem cell niche and are believed to stimulate their differentiation. We hypothesized that if testicular peritubular macrophages do indeed stimulate spermatogonial differentiation, MEHP exposure would result in an increase of peritubular macrophages to stimulate the replacement of lost spermatocytes. Male rats were exposed to 700 mg/kg MEHP or corn oil (vehicle control) *via* oral gavage at PND 28 and euthanized at 48 hours, 1 week, or 2 weeks later. Tubules were stained with immunofluorescent markers for macrophages and undifferentiated spermatogonia. Peritubular macrophages were observed in rat testis similar to those previously described in mice: MHC-II^+^ cells on the surface of seminiferous tubules with heterogeneous morphology. Quantification of MHC-II^+^ cells revealed that, unlike in the mouse, their numbers did not increase through puberty. MEHP increased macrophage presence by six-fold 48-hours after exposure and remained elevated by two-fold two weeks after exposure. An increase of differentiating spermatogonia occurred two weeks after MEHP exposure. Taken together, our results suggest that peritubular macrophages play a crucial role in the testis response to acute injury and the subsequent recovery of spermatogenesis.

**Summary Sentence:** Phthalate-induced testicular injury results in an increase of specialized peritubular macrophages that may assist in the recovery of spermatogenesis.

## Introduction

Di-(2-ethylhexyl) phthalate (DEHP) is a ubiquitous plasticizer that is not chemically bound to its substrates, which results in its release and eventual ingestion by humans and animals [1,2]. Mono-(2-ethylhexyl) phthalate (MEHP) is the primary metabolite of DEHP and has been observed at measurable concentrations in a majority of newborns [3] and in pre-pubertal children from those observed in the US [4] and Germany [5]. MEHP is considered an endocrine disrupting chemical (EDC) that is correlated with reduced free testosterone [6] and increased prolactin and estradiol [7] in adult male humans. MEHP persists in measurable quantities in semen [8] and urine and is correlated with a decrease in sperm quality [9]. Hence, one of the primary concerns regarding MEHP exposure in humans is the deleterious effect it may have on fertility and reproductive function.

Experimental animal models provide a majority of the mechanistic evidence that links phthalate exposure to adverse fertility outcomes. Toxicant exposure, including MEHP, in the testes primarily affect Sertoli cells [10], which sequester meiotic, and therefore antigenic, germ cells behind a selective blood-testis barrier (BTB). The BTB protects germ cells from the immune system, transfer necessary nutrients and information from the endocrine system, and clean up apoptotic germ cells and waste products created during spermatogenesis (Reviewed in: [11]). Acute high dose MEHP exposure in rodents results in the disruption of the Sertoli cell cytoskeleton [12]. A secondary consequence of this disruption is the dysregulation of the apoptosis inducing FAS/FAS-ligand system [13] which results in the widespread death of maturing spermatocytes [14]. In addition, MEHP exposure causes disruption in the gap junctions between Sertoli cells that form the BTB [15] which is, in part, regulated by sTNFα [16], a compound that is capable of initiating an inflammatory immune response.

Indeed, a brief but large monocyte influx appears coincident with widespread spermatocyte apoptosis following MEHP exposure [17]. This influx is at least in part elicited by canonical pro-inflammatory cytokines (e.g. IL-1α, IL-6 and MCP-1) released by Sertoli cells and not by spermatocyte apoptosis itself [18]. Although the infiltrating monocytes observed due to acute MEHP exposure are inflammatory in phenotype, they do not contribute to or exacerbate spermatocyte apoptosis [19].

The purpose or function of the monocyte infiltration into the testes due to MEHP-induced injury remains unknown. Macrophages, which mature from naïve monocytes, make up a large portion of all cells in the testicular interstitium and are the most abundant immune cell type in the testis [20]. Testicular macrophages can be broadly categorized as those resident to the testes and those that infiltrate to the testes as a part of an inflammatory response. Resident testicular macrophages perform a number of essential functions during development and adulthood (Reviewed in: [21,22]). These functions include morphogenesis *via* vascularization during organogenesis [23], assisting in the steroidogenesis of testosterone [24] and, therefore, aiding spermatogenesis and maintaining an immunosuppressive environment [25,26] presumably to protect the integrity of the meiotic spermatogonial niche. Even under inflammatory conditions, resident testicular macrophages have a unique response and tend to polarize towards an anti-inflammatory phenotype; in response to LPS [25], bacteria [27], or IL-4 [28]. Recently, a novel role for testicular macrophages was proposed in mice. A subset of macrophages were observed on the surface of seminiferous tubules, concentrated in areas of undifferentiated spermatogonia, and were found to aid in the differentiation of spermatogonia [29]. This specialized population of macrophages, called peritubular macrophages, were found to be derived exclusively from infiltrating populations of monocytes [30]. Therefore, we hypothesize that the monocyte infiltration observed due to MEHP testicular damage may contribute to the seeding of peritubular macrophages and aid in the recovery of the spermatogonial niche, rather than cause additional damage as previously predicted. To test this hypothesis, we first evaluated if peritubular macrophages are present in the rat testis, then describe their morphological and expression phenotype, and, finally, determine if they associate with the spermatogonial niche.

## Materials And Methods

### Animals and Housing

All animal work was conducted using humane procedures that were preapproved by the Institutional Animal Care and Use Committee at The University of Texas at Austin and followed NIH guidelines (AUP-201900115). Males Fischer CDF344 rats were purchased from Harlan (Indianapolis, IN) and shipped to The University of Texas at Austin at post-natal day (PND) 25. Upon arrival, animals were maintained in a temperature-controlled housing facility on a 12:12 L:D cycle with *ad libitum* access to food and standard rat chow. After 3 days of habituation, on PND 28, males were randomly assigned to one of two treatment exposure groups; vehicle control (VEH) or MEHP.

Each animal was exposed *via* oral gavage to 700 mg/kg MEHP dissolved in corn oil or an equivalent volume of corn oil (VEH). For the data shown in this paper, 7 animals were used in each treatment group in 2 cohorts (3 animals per treatment per age in cohort 1 and 4 animals per treatment per age in cohort 2). Following gavage, animals were checked for general health but were otherwise left undisturbed in their home cage until sacrifice 48 hours after treatment, 1 week after treatment, or 2 weeks after treatment. Animals were euthanized by CO2 asphyxiation followed by cervical dislocation. The testes were rapidly removed and weighed. One testis was randomly selected for whole tubule immunofluorescence and was immediately placed in cold PBS (Gibco, Cat# 10010-023) on ice. The opposite testis was submerged in Bouin’s Fixative on a gentle rocker overnight. Bouin’s was replaced with lithium-saturated ethanol the following day and every subsequent day until the solution and the testis were clear.

Seminiferous tubules were isolated within 1-hr of sacrifice from testes that had been placed in cold PBS on ice. The testis was decapsulated by puncturing and tearing the tunica and the tubules evacuated into cold PBS. The tubules were gently teased apart and the interstitial tissue removed with blunt forceps. The tubules were then washed 4 times in ice cold PBS to further remove interstitial tissue and cells. The tubules were then fixed in 4% paraformaldehyde (Electron Microscopy Sciences, 15710-S) overnight at 4 °C, washed an additional 4 times in cold PBS, and stored in cold PBS at 4 C until use.

Immunofluorescence staining was performed in 96 well plates on seminiferous tubules with primary antibodies major histocompatibility complex class II [MHC Class II RT1D (BioLegend, Clone OX-17, 205401, 1:200)], cluster of differentiation 68 [CD68 (Bio Rad, MCA341R, 1:200)], cluster of differentiation 163 [CD163 (Invitrogen PA5-78961 & MA5-16656, Bio Rad, MCA34212, 1:200)], zinc finger and BTB domain-containing protein 16 [ZBTB16/PLZF (Santa Cruz Biotechnology, sc-22839, 1:200)] and secondary antibodies Alexa 488 (Invitrogen, A11001, 1:500) and Alexa 568 (Invitrogen, A11036, 1:500). Hoescht 33342 (Life Technologies, H3570, 1:10000) was used as a DNA/nuclei counter stain. Stained tubules were mounted whole on super frost slides with Flouromount-G (Southern Biotech, 0100-01) and sealed with standard clear nail polish. Slides were stored at −20 °C until imaging.

Slides were imaged on a Zeiss scanning confocal microscope (Zeiss, LSM 710). Images that were used to count macrophages and spermatogonia were taken at 20x (425.10 μm by 425.10 μm) and with a depth of 15 um in 3 um increments to ensure the surface of the seminiferous epithelium and spermatogonia were captured. Z-stacks were flattened in ImageJ (version 1.49T) using the “Max Intensity” method. Positively stained cells in flattened images were then counted with EBImage (version 4.24.0) and R (version 3.5.3). Cell count was normalized to tubule area and are represented as cell counts per 10^5^ pixel area. An aggregate cell count was calculated across 15 individual tubule observations within each animal by summing the cell count and tubule area of each image and calculating the density of positive cells (# cells/10^5^ area). This methodology considers each animal as the individual statistical unit rather than each tubule observation as the individual statistical unit and avoids violations of non-independence assumptions within statistical tests. Counts were intermittently verified by manual counting in ImageJ to ensure accuracy. Cell counts were compared between treatments and within age using a Wilcoxon Ranked-Sum test because the data were not normally distributed and were considered significant if p < 0.01. Graphs were made in R with ggplot2 (version 3.1.1) and edited only for style in Adobe Illustrator (version CS5). Images for morphological and co-staining analyses were taken at 63x (134.95 μm by 134.95 μm) at z-depths that varied depending on the depth of the cell being imaged.

## Results

Seminiferous tubules from PND30 Fischer CD344 male rats were separated from interstitial tissue, subjected to immunofluorescence for MHCII, and mounted whole on slides for analysis on a confocal microscope. A population of MHCII^+^ cells was observed on the surface of seminiferous tubules (20x – **Fig 1A**). Closer observation of these cells revealed irregular morphology of the cell membrane and nucleus (63x – **Fig1B** and **C**). MHCII^+^ cells were sparse and found on average once every 2 randomly captured (425.1 μm by 425.1 μm, average tubule area 102,084 μm^2^) windows. These cells were located on the surface of tubules often overlying or nestled in-between, in the same plane, or in close proximity to the nuclei of peritubular myoid cells (**Fig 2A & B**), which were identified by their location and unique morphology. Occasionally, long (∼50 - 100 μm) protrusions originating from MHCII^+^ cells appeared to extend around or between the nuclei of peritubular myoid cells (**Fig 2C & D**). In rare instances the nuclei of an MHCII^+^ cell was observed under what appeared to be the nuclei of peritubular myoid cells (**Fig 2E-H**).

**Figure 1.**
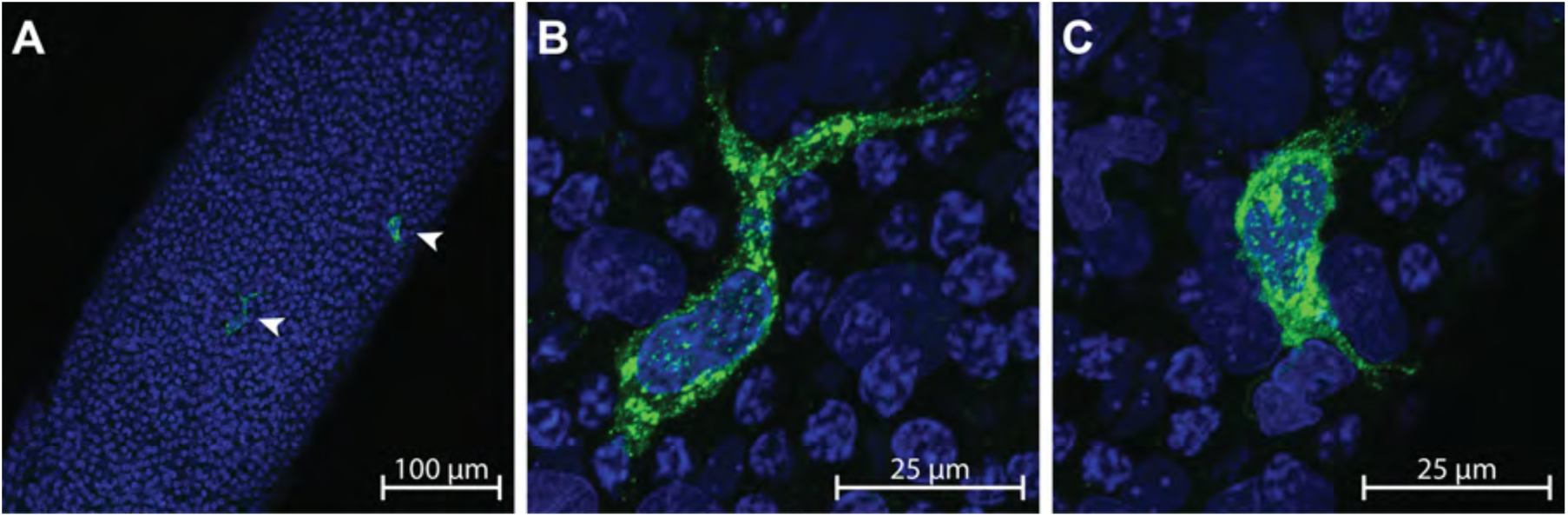
A) MHCII^+^ peritubular macrophages (arrow heads) are shown on the surface of a whole seminiferous tubule at 20x; scale bar = 100 μm. B-C) Up close (63x; scale bar = 25 μm) confocal images show two MHCII^+^ peritubular macrophages with heterogeneous morphology on the surface of a seminiferous tubule.

**Figure 2.**
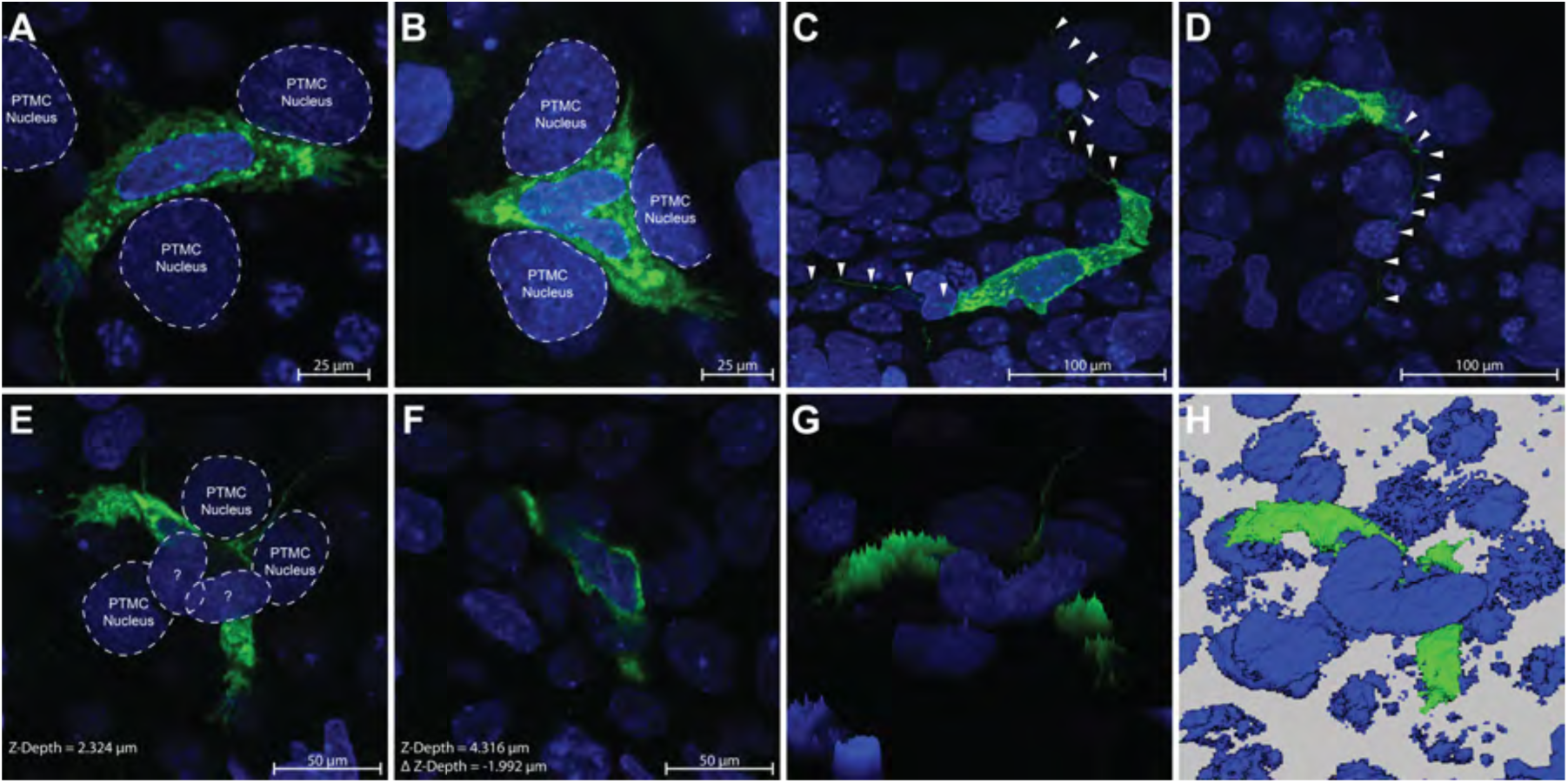
A-B) Two separate macrophages are shown at high magnification (63x; scale bar = 25 μm) nestled between the nuclei of peritubular myoid cells. The macrophage cell surface marker MHCII can clearly be seen to occupy exclusive space in the same plane and between the peritubular myoid cell nuclei. C-D) Long (> 100 μm) extensions can be seen protruding bi-directionally or unidirectionally (respectively) from MHCII^+^ cells, sometimes meandering under the nuclei of other cells (63x; scale bar = 25 μm). E-F) An MHCII^+^ cell is shown in two different scanning planes (E = Z-Depth 2.324 μm and F = Z-Depth 4.316 μm) that is nestled between three peritubular myoid cell nuclei and under the nuclei of two cells that appear to also be peritubular myoid cells (63x; scale bar = 50 μm). G) A 3D brightness histogram and H) a reconstructed 3D image clearly show an MHCII^+^ cell under the nuclei of what appear to be peritubular myoid cells.

The morphology of MHCII^+^ cells was heterogeneous. The majority of MHCII^+^ cells observed on the surface of the seminiferous tubule were either circular (**Fig 3A**) or spindeloid (**Fig 3B --** [31]) in morphology. Observation of MHCII^+^ cells with elongated or stellate morphology was rare, but was occasionally observed (**Fig 3C** and **Fig 3D**, respectively).

**Figure 3.**
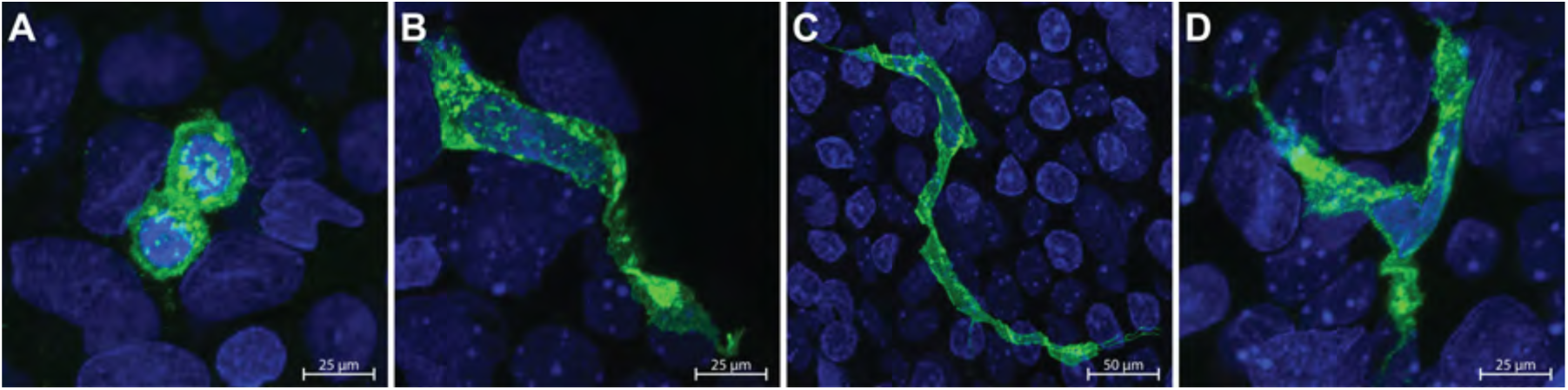
The various morphologies of MHCII^+^ peritubular macrophages observed on the surface of seminiferous tubules is shown; A) circular (63x; scale bar = 25 μm), B) spindeloid (63x; scale bar = 25 μm), C) elongated (63x; scale bar = 50 μm), D) stellate (63x; scale bar = 25 μm).

In order to determine if the MHCII^+^ cells observed were in fact derived from the monocyte lineage, tubules were double stained with MHCII^+^ and the lysosomal protein CD68. All observed MHCII^+^ cells on the surface of seminiferous tubules were CD68^+^ regardless of the morphology of MHCII^+^ cells (**Fig 4**). All MHCII^+^ cells remained CD68^+^ regardless of the age or treatment of the animal analyzed (PND30; **Fig 4A-B**, PND 35; **Fig 4C-D**, PND 42; **Fig 4E-F**).

**Figure 4.**
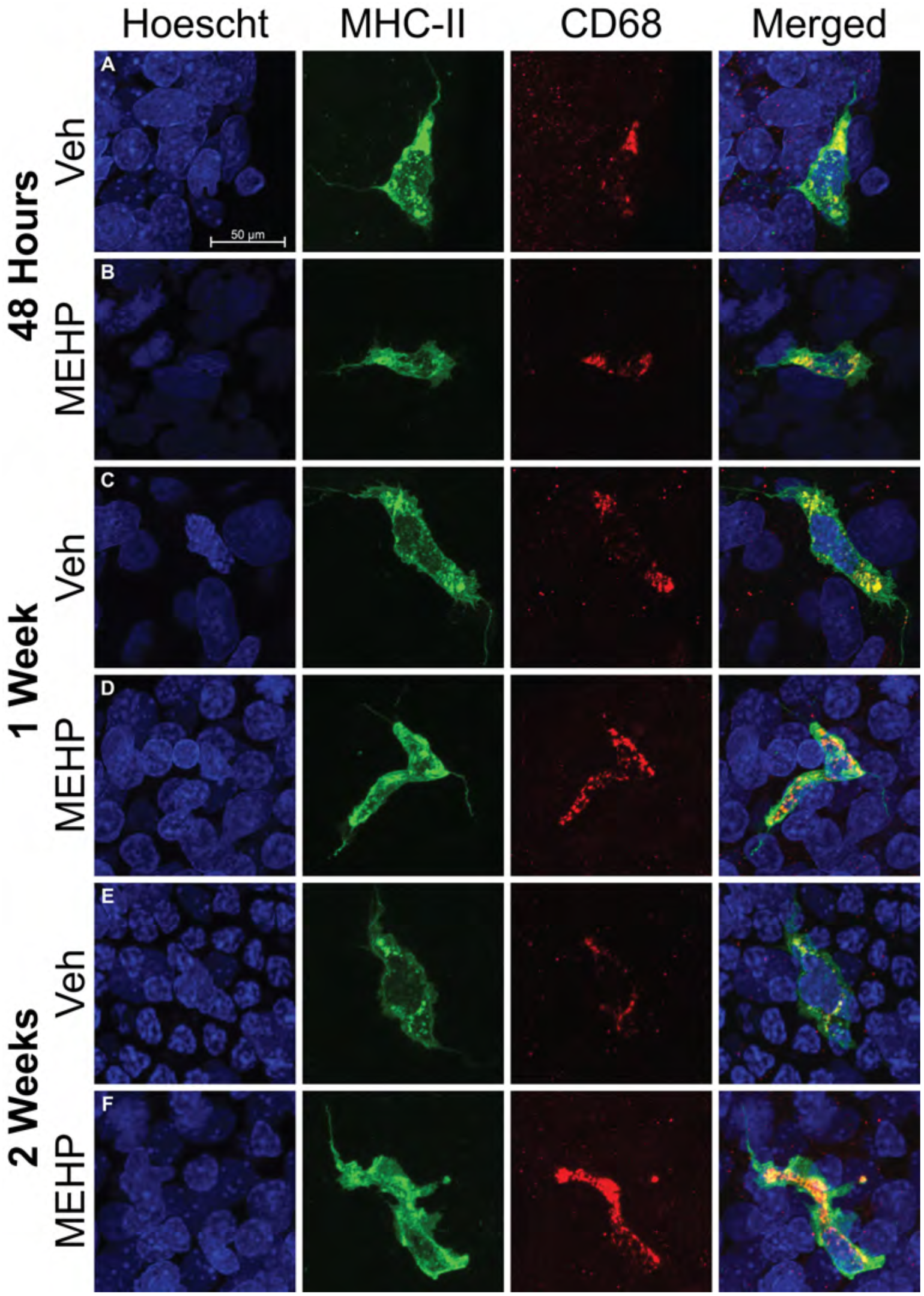
Dual staining of peritubular macrophages with MHCII (green) and CD68 (red) are shown at all treatment and age endpoints; A-B) 48 hours after MEHP exposure, C-D) 1 week after MEHP exposure, and E-F) 2 weeks after MEHP exposure. The Hoescht (blue) nuclei counter stain is used in all images and the scale bar shown in part A is the same for all images. All MHCII^+^ macrophages are also CD68^+^ regardless of treatment, age, or time after exposure. All imaged were taken at 63x. The scale bar (50 μm) represented in panel A applies to all images in this figure.

We hypothesized that if the function of peritubular macrophages is indeed to stimulate the differentiation of spermatogonial cells, then depletion of maturing germ cells via MEHP treatment would induce the mobilization of peritubular macrophages to the seminiferous tubules. PND 28 male rats were exposed to either MEHP or corn oil (VEH) and sacrificed after 48 hours, 1 week, or 2 weeks with no further manipulation and the number of MHCII^+^ cells, controlled for by tubule area, was counted (n = 7 per treatment per age – **Fig 5**). After 48 hours, the population of MHCII^+^ was significantly elevated ∼ 6.5-fold compared to VEH males of the same age (U = 3, p = 0.002 – **Fig 6**). The number of MHCII^+^ cells began to decline but remained significantly elevated in MEHP animals ∼ 6 fold after 1 week (U = 0, p = 0.0003) and ∼ 2.5-fold after 2 weeks (U = 6, p = 0.005). Age did not affect the number of MHCII^+^ cells in VEH animals as determined by a Kruskal-Wallis test (H(2) = 4.1039, p = 0.1285). Subsequent pairwise Kruskal-Wallis rank sum tests, adjusted with the Benjamini-Hochberg method, also failed to identify any differences in MHCII^+^ cell counts across age and within VEH animals.

**Figure 5.**
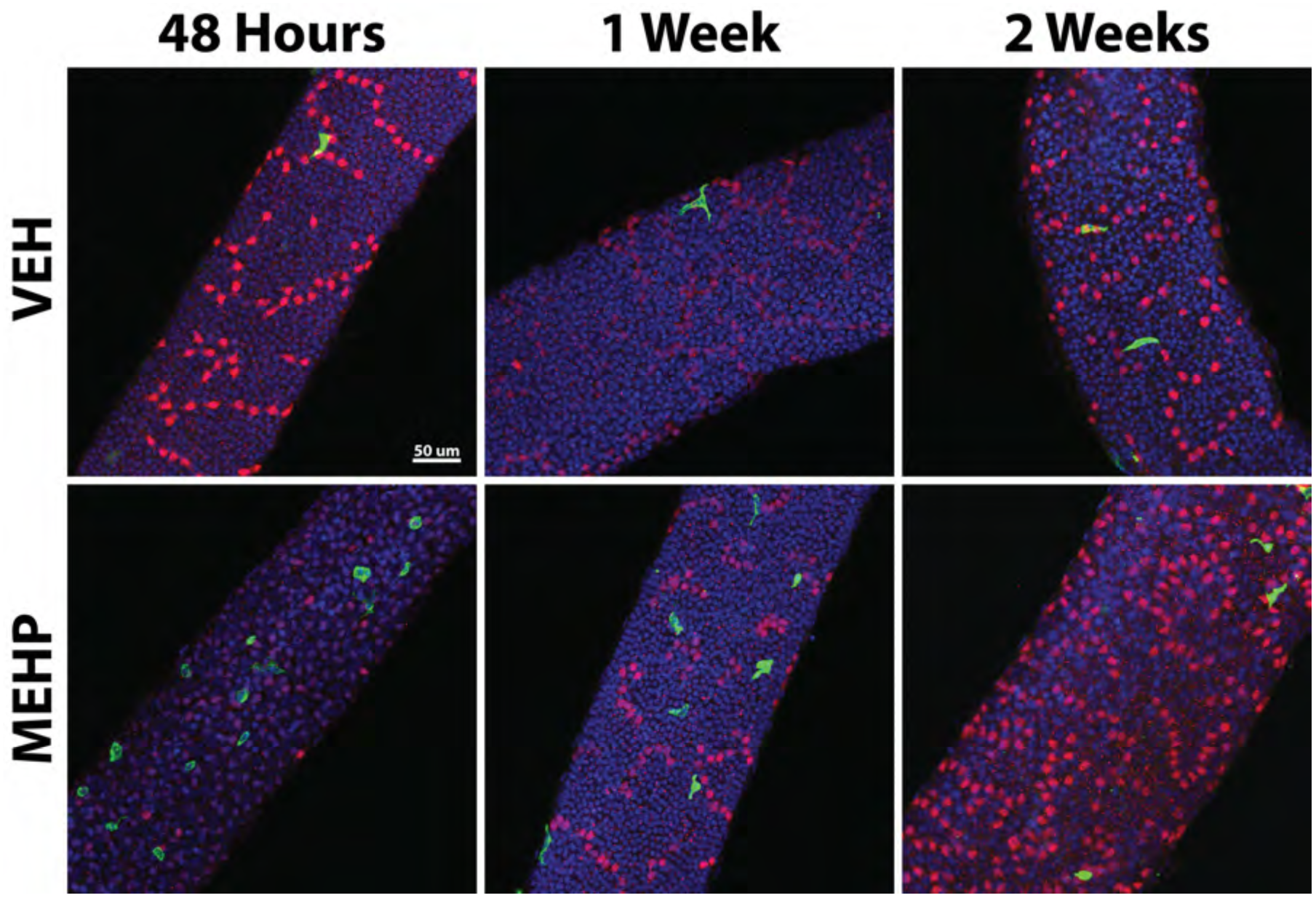
Representative micrographs of MHCII^+^ (green) peritubular macrophages are shown on the surface of seminiferous tubules in close proximity to PLZF^+^ (red) spermatogonia in both VEH and MEHP treated rats. At 48 hours after exposure, dense clusters of primarily circular MHCII^+^ cells are observed on tubules with more single PLZF^+^ cells. Circular MHCII^+^ cells are not seen in VEH treated or 1 week/2 week MEHP treated samples and the maximum density of MHCII^+^ cells decreases at 1week/2 weeks after MEHP exposure. All images in this figure were taken at 20x. The scale bar (50 μm) represented in the top left panel applies to all images in this figure.

**Figure 6.**
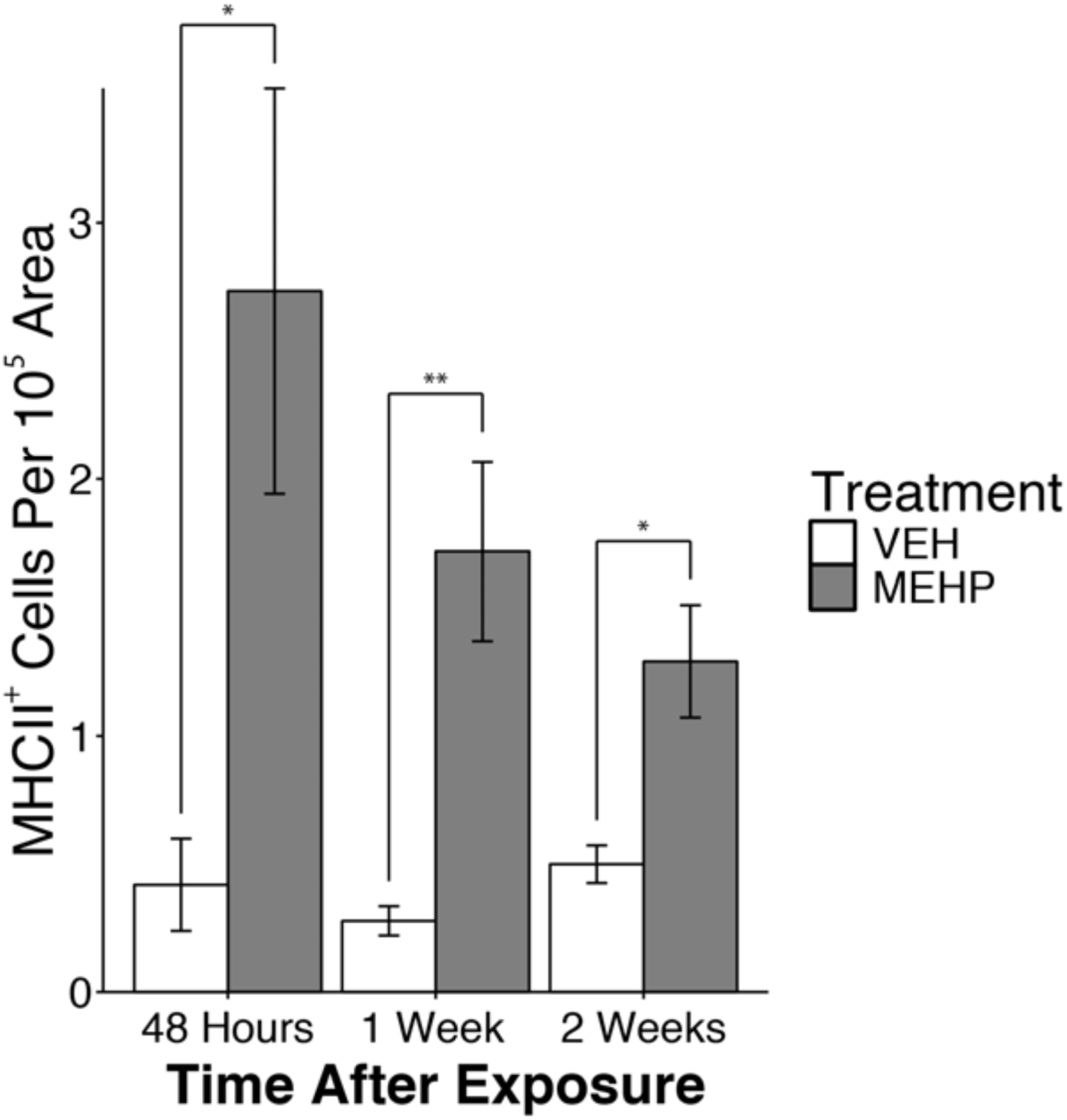
The number of MHCII^+^ cells, normalized to tubule area, is shown. In VEH treated males, the number of peritubular macrophages does not increase through puberty as previously reported in mice, but stays at a constant low level. At all time points, the number of MHCII^+^ cells on the surface of seminiferous tubules is elevated in males exposed to MEHP; * p < 0.01 and ** p < 0.001.

In order to determine the distribution of MHCII^+^ cells, a density analysis of the number of MHCII^+^ cells per tubule area was performed (**Fig 7**). Males exposed to the VEH treatment, which represent a “normal” developmental trajectory, at the 48 hour and 1 week time point had a large proportion of instances (∼75%) that had no MHCII^+^ cells. The number of instances with zero MHCII^+^ cells was reduced to 56% in VEH treated males at 2 week time point, which suggests that while the total number of MHCII^+^ doesn’t change with age, their localization does. The distribution of MHCII^+^ cells in MEHP treated males was markedly different. Instances with no MHCII^+^ cells was only 32% at 48 hours and 2 weeks and was slightly decreased at the 1 week time point (25%). There were isolated observations of high-density concentrations of MHCII^+^ cells in MEHP treated animals (>8 MHCII^+^ cells per 10^5^ area – 12% at 48hrs and 1% at 1 week) but none in the VEH treated groups. The majority of the increase in MHCII^+^ cells seen due to MEHP treatment was observed in intermediate densities (< 8 MHCII^+^ cells per 10^5^ area).

**Figure 7.**
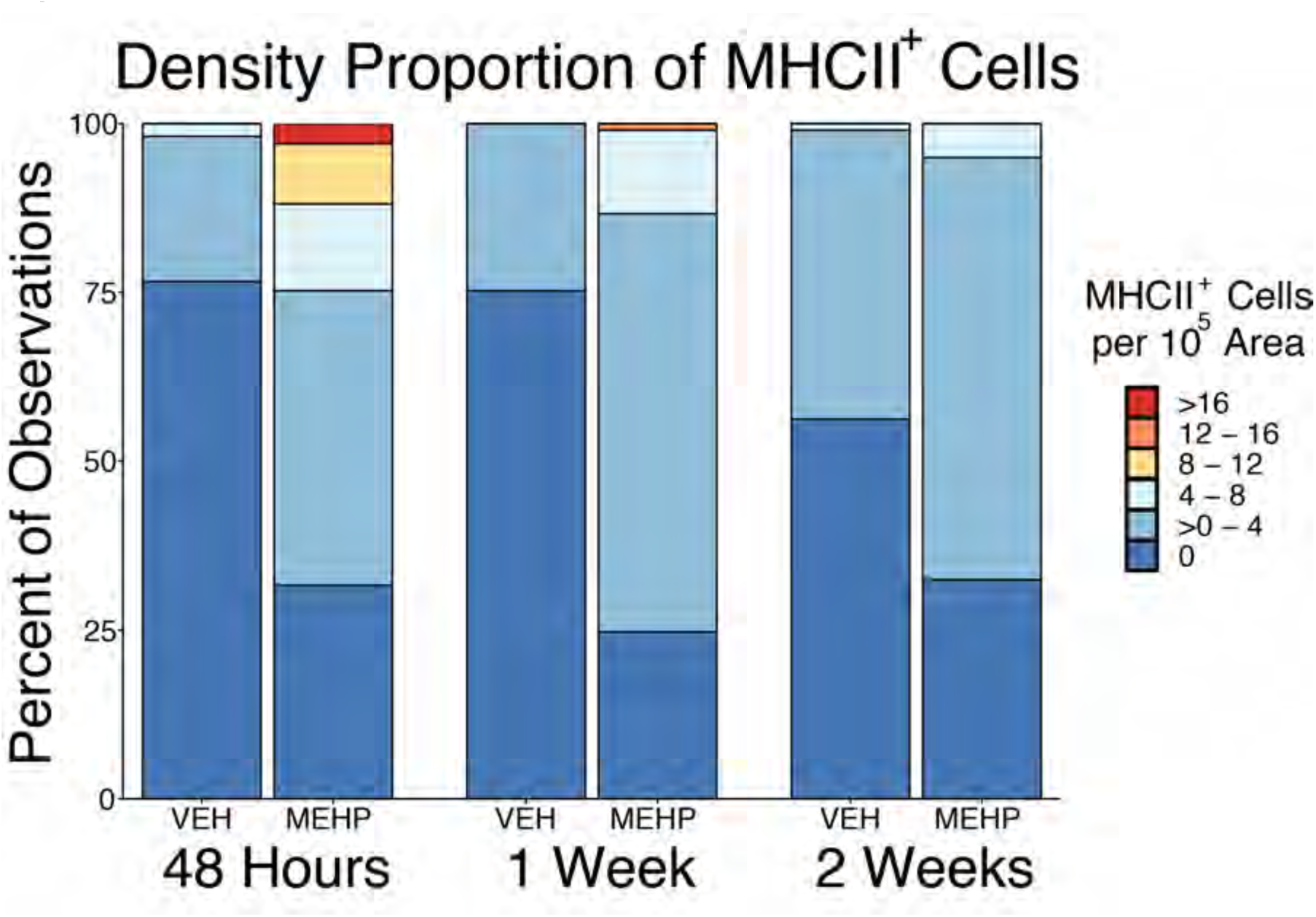
The proportion of individual observations as a function of density (# MHCII^+^ cells/10^5^ tubule area) is shown. Observations in VEH treated animals show a majority (∼75%) of instances with no MHCII^+^ cells at 48 hours and 1 week after treatment, which is diminished at 2 weeks. In MEHP treated males, the proportion of tubule observations with no MHCII^+^ cells is lower (∼30-25%). Although high densities of MHCII^+^ cells were observed in MEHP treated males, the majority of the observed increase was due to moderate densities.

Finally, we stained whole tubules with PLZF to determine if spermatogonial number was affected by MEHP treatment. We found no difference in the number of PLZF^+^ cells at either the 48-hour or 1 week time point. However, the number of PLZF^+^ cells was increased due to MEHP exposure at the 2-week time point (U = 3, p < 0.001 – **Fig 8**).

**Figure 8.**
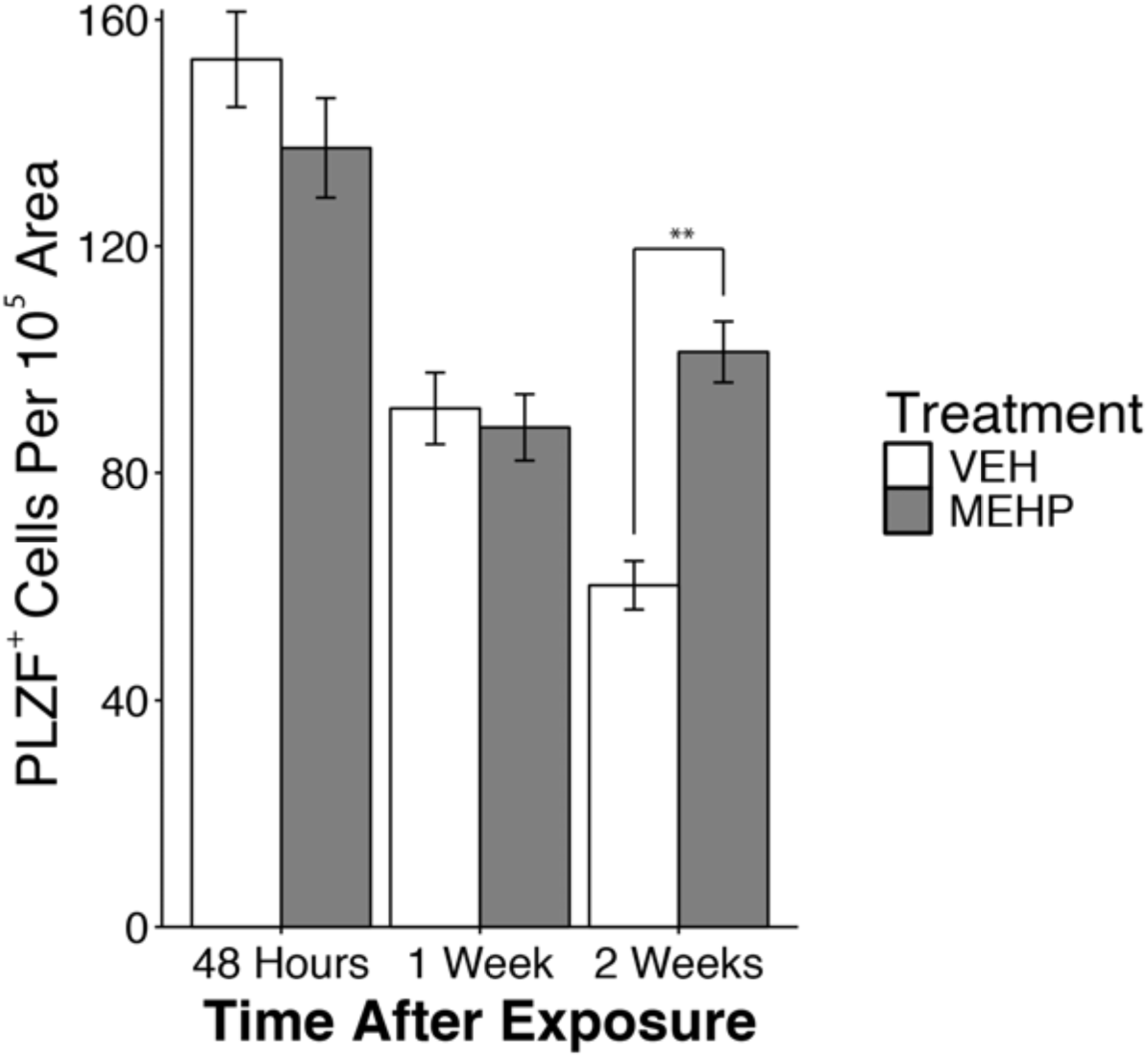
The number of PLZF^+^ spermatogonia in VEH and MEHP treated male rats is shown normalized to tubule area. The PLZF^+^ count appears to diminish with age, but is primarily a factor of area normalization, which is attributed to the expansion of tubules as spermatogenesis progresses through puberty. MEHP treated males show an increase (** p < 0.001) of PLZF^+^ spermatogonia 2 weeks after exposure.

## Discussion

Testicular macrophages perform an eclectic set of functions in the testes that span, and are essential to, the life cycle (Reviewed in: [22]). During development, testicular macrophages aid in the organogenesis of the testis by directing vascularization [23]. During adolescence, testicular macrophages stimulate the differentiation and proliferation of Leydig cells, the prime steroid-hormone producing cell in the testis [32]. In adulthood, testicular macrophages aid in the ultimate role of the testes, spermatogenesis, by aiding in the steroidogenesis of testosterone [24]. Throughout the lifespan, testicular macrophages help maintain an immune-privileged environment in the testis and therefore they protect sanctity of spermatogenesis [33]. Recently, a newly discovered function was ascribed to testicular macrophages. A subset of testicular macrophages (peritubular macrophages) was identified in mice that localize to the surface of seminiferous tubules near the spermatogonial niche that when depleted disrupts spermatogonial differentiation [29].

The current work aimed to describe the peritubular subset of testicular macrophages in the rat for the first time, characterize their morphology and distribution, and observe their newly ascribed function in an injury model. While previous models have shown that depleting macrophages from the testis reduces the number of differentiating spermatogonia [29], here we demonstrate that depletion of spermatocytes by exposure to MEHP mobilizes peritubular macrophages to the surface of seminiferous tubules in large numbers for at least 2 weeks. This increase is concurrent with an increase in the number of differentiating spermatogonia. This work is significant in that it is one of the first demonstrations that peritubular macrophages are specifically mobilized by testicular injury and the large-scale loss of spermatocytes.

MEHP is a well described Sertoli cell toxicant [12], which causes widespread loss of spermatocytes [14]. The loss of spermatocytes due to MEHP exposure is age- (younger > older), species- (rat > mouse), and dose-dependent [17]. The loss of spermatocytes is concurrent with a massive but transient (∼48 hours) monocyte influx that is congruent with the degree of spermatocyte loss; the monocyte influx is more robust in younger animals, larger in rats compared to mice and larger at higher doses of MEHP [17]. It was initially hypothesized that because the monocyte influx induced by MEHP was proportional to the loss of spermatocytes, this large-scale inflammatory immune response might exacerbate, or be the cause of, the loss of spermatocytes and thus account for the specificity of the observed effects. This was found not to be the case. The depletion of circulating monocytes, which neutralizes the MEHP induced monocyte influx, has no effect on spermatocyte apoptosis [19]. The precipitating cause and the functional purpose of this influx is not yet fully understood, but our current results suggest that at least one consequence is an increase in the long-term (2 week) presence of peritubular macrophages.

Given that the only known function of peritubular macrophages is to stimulate the differentiation of spermatogonia, and that a loss of spermatocytes would necessitate their replacement by the induction of spermatogonial differentiation, it is reasonable to suggest that the increase in peritubular macrophages we observe is to stimulate spermatogonial differentiation to replace the lost spermatocytes. Our data indirectly support this hypothesis; 2 weeks after MEHP exposure, a time point where peritubular macrophages remain elevated, the number of PLZF^+^ cells (differentiating spermatogonia) is increased. This hypothesis requires additional evidence and experiments, which are currently underway in our laboratory.

Peritubular macrophages may, of course, have functions other than, or in addition to, the stimulation of spermatogonial differentiation. There are two primary immunoreactivity profiles of macrophages that populate the testes in rats; tissue resident testicular-macrophages (CD68^-^ /CD163^+^) and newly arrived monocyte-like testicular macrophages (CD68^+^/CD163^-^) (Reviewed in: [21]). These two populations have distinct expression phenotypes that represent antiinflammatory tissue resident macrophages (CD68^-^/CD163^+^) and “alternatively activated macrophages” with suppressed inflammatory activity (CD68^+^/CD163^-^) [28].

We found that all peritubular macrophages in rats are CD68^+^, indicating that they are either derived from circulating monocytes that quickly (within 48 hours) transition and localize to the seminiferous tubules or they constantly maintain the CD68^+^ phenotype regardless of when they arrived in the testes. It is noteworthy that all peritubular macrophages in VEH animals were also CD68^+^ at all time points, which supports the latter possibility; they maintain a CD68^+^ phenotype. This is consistent with previous observations in mice that found macrophages are exclusively seeded from bone-marrow derived circulating monocytes using lineage-tracing techniques [30]. Determining a definitive immunoreactive profile for peritubular macrophages in future experiments is essential so that further characterization can be performed (e.g. fluorescent activated cell sorting to allow for macrophage separation and transcriptome profiling). Efforts are currently underway to definitively describe the CD163 phenotype of peritubular macrophages in rats.

The location and morphology of peritubular macrophages also provide some evidence of their possible function. Sertoli cells can perform macrophage-like functions and therefore serve an immune cell role in the seminiferous epithelium; phagocytosis and ingestion of apoptotic spermatocytes [34] and microbe detection via the Toll-like receptor [35]. Sertoli cells actively participate in the clearance of apoptotic spermatocytes [36] and secrete the powerful monocyte chemoattractant protein-1 (MCP-1) [18] specifically due to MEHP induced injury. Under normal physiological conditions, Sertoli cells can induce germ cell apoptosis and ingest said cells such that the meiotic and immunogenic remnants remain sequestered behind the blood-testis-barrier (Reviewed in: [37]). It is, however, possible that in extreme cases of spermatocyte loss, such as those induced by acute MEHP exposure, Sertoli cells are not capable of keeping up with the processing of apoptotic bodies. A possible consequence of this scenario is that if the blood-testis-barrier becomes compromised, an effect observed due to MEHP exposure [15], an auto-antigenic immune response could be mounted against the “non-self” antigens and cause further damage. Sertoli cells can act as antigen presenting cells which diminishes inflammatory immune responses [38]. That peritubular macrophages reside in such close proximity to Sertoli cells, that they are themselves antigen presenting cells, and that they respond specifically to an event with widespread apoptosis, suggests that they may play a role in the tolerization of inflammatory immune responses in certain catastrophic scenarios. Direct macrophage cell-cell contact and communication in the testis is not unheard of. Interstitial macrophages are known to directly communicate with Leydig cells and aid in steroidogenesis [39]. That peritubular macrophages could contact and interact with Sertoli cells would not be unlikely.

The work presented here demonstrates that a specialized sub-category of testicular macrophage infiltrates and occupies the testes due to acute testicular injury in rats. This work is fundamental to understanding the mechanisms by which infertility and azoospermia occur in adult males. Correlational observations in humans consistently note an increased presence of CD68^+^ macrophages in infertile men [40,41]. These studies suggest the presence of CD68^+^ macrophages may contribute to or be the cause of infertility. These studies, however, are retrospective in nature and may misinterpret the significance of the increased presence of macrophages. Although more experimentation is needed, our data, and that of our predecessors, suggests instead that CD68^+^ peritubular macrophages initially infiltrate the testis because of the loss of maturing spermatocytes and to stimulate spermatogonial differentiation.

## References

[1] Takehisa H, Naoko E, Masahiko S, Katsuhide T, Moriyuki O, Keizoh S, Mutsuko T, Kenji K, Shin’ichiro N, Toshio O. Release behavior of diethylhexyl phthalate from the polyvinyl-chloride tubing used for intravenous administration and the plasticized PVC membrane. International Journal of Pharmaceutics 2005; 297:30–37.

[2] Hanawa T, Muramatsu E, Asakawa K, Suzuki M, Tanaka M, Kawano K, Seki T, Juni K, Nakajima S. Investigation of the release behavior of diethylhexyl phthalate from the polyvinyl-chloride tubing for intravenous administration. International Journal of Pharmaceutics 2000; 210:109–115.

[3] Latini G, De Felice C, Presta G, Del Vecchio A, Paris I, Ruggieri F, Mazzeo P. In utero exposure to di-(2-ethylhexyl)phthalate and duration of human pregnancy. Environ Health Perspect 2003; 111:1783–1785.

[4] Silva MJ, Barr DB, Reidy JA, Malek NA, Hodge CC, Caudill SP, Brock JW, Needham LL, Calafat AM. Urinary levels of seven phthalate metabolites in the U.S. population from the National Health and Nutrition Examination Survey (NHANES) 1999-2000. Environ Health Perspect 2004; 112:331–338.

[5] Becker K, Seiwert M, Angerer J, Heger W, Koch HM, Nagorka R, Roßkamp E, Schlüter C, Seifert B, Ullrich D. DEHP metabolites in urine of children and DEHP in house dust. Int J Hyg Environ Health 2004; 207:409–417.

[6] Pan G, Hanaoka T, Yoshimura M, Zhang S, Wang P, Tsukino H, Inoue K, Nakazawa H, Tsugane S, Takahashi K. Decreased serum free testosterone in workers exposed to high levels of di-n-butyl phthalate (DBP) and di-2-ethylhexyl phthalate (DEHP): a cross-sectional study in China. Environ Health Perspect 2006; 114:1643–1648.

[7] Li S, Dai J, Zhang L, Zhang J, Zhang Z, Chen B. An association of elevated serum prolactin with phthalate exposure in adult men. Biomed Environ Sci 2011; 24:31–39.

[8] Han SW, Lee H, Han SY, Lim DS, Jung KK, Kwack SJ, Kim KB, Lee BM. An exposure assessment of di-(2-ethylhexyl) phthalate (DEHP) and di-n-butyl phthalate (DBP) in human semen. J Toxicol Environ Health Part A 2009; 72:1463–1469.

[9] Wang Y-X, You L, Zeng Q, Sun Y, Huang Y-H, Wang C, Wang P, Cao W-C, Yang P, Li Y-F, Lu W-Q. Phthalate exposure and human semen quality: Results from an infertility clinic in China. Environ Res 2015; 142:1–9.

[10] Boekelheide K, Neely MD, Sioussat TM. The Sertoli cell cytoskeleton: a target for toxicant-induced germ cell loss. Toxicol Appl Pharmacol 1989; 101:373–389.

[11] França LR, Hess RA, Dufour JM, Hofmann MC, Griswold MD. The Sertoli cell: one hundred fifty years of beauty and plasticity. vol. 4. John Wiley & Sons, Ltd (10.1111); 2016.

[12] Richburg JH, Boekelheide K. Mono-(2-ethylhexyl) phthalate rapidly alters both Sertoli cell vimentin filaments and germ cell apoptosis in young rat testes. Toxicol Appl Pharmacol 1996; 137:42–50.

[13] Lee J, Richburg JH, Shipp EB, Meistrich ML, Boekelheide K. The Fas system, a regulator of testicular germ cell apoptosis, is differentially up-regulated in Sertoli cell versus germ cell injury of the testis. Endocrinology 1999; 140:852–858.

[14] Richburg JH, Nañez A, Gao H. Participation of the Fas-signaling system in the initiation of germ cell apoptosis in young rat testes after exposure to mono-(2-ethylhexyl) phthalate. Toxicol Appl Pharmacol 1999; 160:271–278.

[15] Yao P-L, Lin Y-C, Richburg JH. Mono-(2-ethylhexyl) phthalate-induced disruption of junctional complexes in the seminiferous epithelium of the rodent testis is mediated by MMP2. Biology of Reproduction 2010; 82:516–527.

[16] Yao P-L, Lin Y-C, Sawhney P, Richburg JH. Transcriptional regulation of FasL expression and participation of sTNF-alpha in response to sertoli cell injury. J Biol Chem 2007; 282:5420–5431.

[17] Murphy CJ, Stermer AR, Richburg JH. Age- and Species-Dependent Infiltration of Macrophages into the Testis of Rats and Mice Exposed to Mono-(2-Ethylhexyl) Phthalate (MEHP)1. Biology of Reproduction 2014; 91:247–11.

[18] Stermer AR, Murphy CJ, Ghaffari R, Di Bona KR, Voss JJ, Richburg JH. Mono-(2-ethylhexyl) phthalate-induced Sertoli cell injury stimulates the production of pro-inflammatory cytokines in Fischer 344 rats. Reproductive Toxicology 2017; 69:150–158.

[19] Voss JJLP, Stermer AR, Ghaffari R, Tiwary R, Richburg JH. MEHP-induced rat testicular inflammation does not exacerbate germ cell apoptosis. Reproduction 2018; 156:35–46.

[20] Niemi M, Sharpe RM, Brown WR. Macrophages in the interstitial tissue of the rat testis. Cell Tissue Res 1986; 243:337–344.

[21] Hedger MP. Macrophages and the immune responsiveness of the testis. J Reprod Immunol 2002; 57:19–34.

[22] Bhushan S, Meinhardt A. The macrophages in testis function. J Reprod Immunol 2017; 119:107–112.

[23] DeFalco T, Bhattacharya I, Williams AV, Sams DM, Capel B. Yolk-sac-derived macrophages regulate fetal testis vascularization and morphogenesis. Proc Natl Acad Sci USa 2014; 111:E2384–93.

[24] Hutson JC. Development of cytoplasmic digitations between Leydig cells and testicular macrophages of the rat. Cell Tissue Res 1992; 267:385–389.

[25] Kern S, Robertson SA, Mau VJ, Maddocks S. Cytokine secretion by macrophages in the rat testis. Biology of Reproduction 1995; 53:1407–1416.

[26] Hayes R, Chalmers SA, Nikolic-Paterson DJ, Atkins RC, Hedger MP. Secretion of bioactive interleukin 1 by rat testicular macrophages in vitro. J Androl 1996; 17:41–49.

[27] Bhushan S, Hossain H, Lu Y, Geisler A, Tchatalbachev S, Mikulski Z, Schuler G, Klug J, Pilatz A, Wagenlehner F, Chakraborty T, Meinhardt A. Uropathogenic E. coli induce different immune response in testicular and peritoneal macrophages: implications for testicular immune privilege. PLoS ONE 2011; 6:e28452.

[28] Winnall WR, Muir JA, Hedger MP. Rat resident testicular macrophages have an alternatively activated phenotype and constitutively produce interleukin-10 in vitro. J Leukoc Biol 2011; 90:133–143.

[29] DeFalco T, Potter SJ, Williams AV, Waller B, Kan MJ, Capel B. Macrophages Contribute to the Spermatogonial Niche in the Adult Testis. Cell Rep 2015; 12:1107–1119.

[30] Mossadegh-Keller N, Gentek R, Gimenez G, Bigot S, Mailfert S, Sieweke MH. Developmental origin and maintenance of distinct testicular macrophage populations. J Exp Med 2017; 214:2829–2841.

[31] Heinrich F, Lehmbecker A, Raddatz BB, Kegler K, Tipold A, Stein VM, Kalkuhl A, Deschl U, Baumgärtner W, Ulrich R, Spitzbarth I. Morphologic, phenotypic, and transcriptomic characterization of classically and alternatively activated canine blood-derived macrophages in vitro. PLoS ONE 2017; 12:e0183572.

[32] Gaytan F, Bellido C, Aguilar E, van Rooijen N. Requirement for testicular macrophages in Leydig cell proliferation and differentiation during prepubertal development in rats. J Reprod Fertil 1994; 102:393–399.

[33] Kern S, Maddocks S. Indomethacin blocks the immunosuppressive activity of rat testicular macrophages cultured in vitro. J Reprod Immunol 1995; 28:189–201.

[34] Nakanishi Y, Shiratsuchi A. Phagocytic removal of apoptotic spermatogenic cells by Sertoli cells: mechanisms and consequences. Biol Pharm Bull 2004; 27:13–16.

[35] Riccioli A, Starace D, Galli R, Fuso A, Scarpa S, Palombi F, De Cesaris P, Ziparo E, Filippini A. Sertoli cells initiate testicular innate immune responses through TLR activation. JI 2006; 177:7122–7130.

[36] Tay TW, Andriana BB, Ishii M, Choi EK, Zhu XB, Alam MS, Tsunekawa N, Kanai Y, Kurohmaru M. Phagocytosis plays an important role in clearing dead cells caused by mono(2-ethylhexyl) phthalate administration. Tissue Cell 2007; 39:241–246.

[37] Print CG, Loveland KL. Germ cell suicide: new insights into apoptosis during spermatogenesis. Bioessays 2000; 22:423–430.

[38] Dal Secco V, Riccioli A, Padula F, Ziparo E, Filippini A. Mouse Sertoli cells display phenotypical and functional traits of antigen-presenting cells in response to interferon gamma. Biology of Reproduction 2008; 78:234–242.

[39] Hutson JC. Changes in the Concentration and Size of Testicular Macrophages during Development. Biology of Reproduction 1990; 43:885–890.

[40] Goluža T, Boscanin A, Cvetko J, Kozina V, Kosović M, Bernat MM, Kasum M, Kaštelan Ž, Ježek D. Macrophages and Leydig cells in testicular biopsies of azoospermic men. Biomed Res Int 2014; 2014:828697.

[41] Frungieri MB, Calandra RS, Lustig L, Meineke V, Köhn FM, Vogt HJ, Mayerhofer A. Number, distribution pattern, and identification of macrophages in the testes of infertile men. Fertil Steril 2002; 78:298–306.

